# Validation and Performance Comparison of Three SARS-CoV-2 Antibody Assays

**DOI:** 10.1101/2020.05.29.124776

**Authors:** Kimberly J Paiva, Ricky D Grisson, Philip A Chan, John R. Lonks, Ewa King, Richard C Huard, Diane L Pytel-Parenteau, Ga Hie Nam, Evgeny Yakirevich, Shaolei Lu

## Abstract

Serology testing of severe acute respiratory syndrome coronavirus 2 (SARS-CoV-2) is increasingly being used during the current pandemic of Coronavirus Disease 2019 (COVID-19). The clinical and epidemiologic utilities of antibody-based SARS-CoV-2 testing are under debate. Characterizing these assays helps to understand the disease and provides scientific basis for deciding how to best use these assays. The study assessed one chemiluminescent assay (Abbott COVID-2 IgG) and two lateral flow assays (STANDARD Q [SQ] IgM/IgG Duo and Wondfo Total Antibody Test). Validation included 113 blood samples from 71 PCR-confirmed COVID-19 patients and 1182 samples from negative controls with potential interferences/cross-reactions, including 1063 pre-pandemic samples. IgM antibodies against SARS-CoV-2 were detected as early as post-symptom onset days 3-4. IgG antibodies were first detected post-onset days 5-6 by SQ assays. The detection rates increased gradually, and SQ IgG, Abbott IgG and Wondfo Total detected antibodies from all the PCR-confirmed patients 14 days after symptom onset. Overall agreements between SQ IgM/IgG and Wondfo Total was 88.5% and between SQ IgG and Abbott IgG was 94.6% (Kappa = 0.75, 0.89). No cross-reaction with other endemic coronavirus infections were identified. Viral hepatitis and autoimmune samples were the main cross-reactions observed. However, the interferences/cross-reactions were low. The specificities were 100% for SQ IgG and Wondfo Total and 99.62% for Abbott IgG and 98.87% for SQ IgM. These findings demonstrate high sensitivity and specificity of appropriately validated antibody-based SARS-CoV-2 assays with implications for clinical use and epidemiological seroprevalence studies.

## Introduction

There is an ongoing worldwide pandemic caused by a novel coronavirus, now known as SARS-CoV-2. The virus was first reported in Wuhan, Hubei Province of China in 2019. Coronavirus Disease 2019 (COVID-19), the disease caused by SARS-CoV-2, has greatly impacted many countries, most especially the United States. There are currently over 5 million confirmed cases worldwide, with over 1.6 million patients in the United States (https://coronavirus.jhu.edu/map.html). Evaluating the spread and transmission of SARS-Cov-2 is critical in addressing the pandemic.

Development of diagnostic methods for COVID-19 started with SARS-CoV-2 viral genome sequencing first shared by a group of Chinese scientists (1). Real-time reverse transcriptase-polymerase chain reaction (rRT-PCR) based methods have been the mainstay as a diagnostic approach. Most tests use the nasopharyngeal and/or oropharyngeal swabs to obtain the virus before running rRT-PCR. However, it was quickly discovered that the detection rates of pharyngeal and nasal swabs were only 32% and 63%, respectively, and their detection rates decreased as the disease progressed (2). First-time positive rate by pharyngeal swab rRT-PCR was reported as low as 37% in 610 hospitalized patients (3).

While rRT-PCR-based testing is the main tool for clinical diagnosis, antibody-based testing has gained considerable attention. Studies suggested that IgM antibody might develop as early as five days after onset of symptoms (4) and IgG developed later at a median time of 14 days (4, 5). The sensitivities of these tests reportedly varies widely from as low as 11% early in infection (6) to as high as 100% after 14 days (5). It has been shown that diagnosis of COVID-19 could potentially be improved by using both PCR-based and antibody-based tests (5). However, the most important use of antibody-based tests is seroprevalence studies for use in modeling methods and understanding how SARS-CoV-2 has spread across different populations.

Many antibody-based SARS-CoV-2 tests are currently available or in development. Antibodies to the Spike protein (S-protein), receptor binding domain (in S-protein) and nucleocapsid protein (N-protein) are the main targets of these assays. It has been demonstrated that N-protein-based antibody tests were more sensitive than antibody tests targeting S-proteins (7). The majority of the assays on the markets are immunochromatographic assays using a lateral flow format. Lateral flow assays use venous blood or capillary blood and they are a manual test that are quick and easy to perform, independent of larger immunochemical instruments. The majority of tests detect IgM and IgG separately while some detect total antibodies (IgM and IgG). Positive results demonstrate a visible band with various degrees of intensity in a designated zone. Chemiluminescent tests are considered the most sensitive by methodology and provided results with great accuracy and precision. These tests are commonly quick and randomly accessible on immunochemical analyzers. The current study looks to the performance of two lateral flow assays and one chemiluminescent assay testing for SARS-CoV-2 antibodies.

## Methods

The study was approved by the Institutional Review Board (IRB) of Lifespan Health System (including Rhode Island Hospital and The Miriam Hospital) to ensure the study met the ethical requirements.

### Patients

A total of 113 remnant/discarded serum or plasma samples were collected from March to April in 2020 from the Clinical Immunology Lab at a major academic pathology department in Rhode Island. These samples were collected from 71 COVID-19 patients confirmed by rRT-PCR tests on nasopharyngeal swabs. An additional 126 samples were collected from healthy individuals in early March. 119 samples that were positive for antibodies against viruses and other pathogens were used to test cross-reaction of the assays (Table3). Additional samples were collected consisting of interference antibodies such as Rheumatoid factor (RF), anti-double strand DNA (ds-DNA), anti-nuclear antibody (ANA) and paraprotein IgM and IgG (Table 3). Blood samples from patients testing positive for upper respiratory viruses were obtained when a viral respiratory pathogen nucleic acid test was performed (ePlex Respiratory Pathogen Panel, GenMark, Carlsbad, CA), or up to 53 days after the diagnoses. The same upper respiratory virus tests were routinely ordered for all the COVID-19 patients. The tests were performed following manufacturer’s protocol.

**Table 1.** Clinicopathologic parameters of the COVID-19 patients.

| Patient cases | Total | Male | Female | P-value |
| --- | --- | --- | --- | --- |
| Sex | 71 | 42 (59%) | 29 (41%) |  |
| Age (Mean ± SEM) | 59.5 ± 1.9 | 56.2 ± 1.9 | 64.3 ± 3.4 | 0.0470 |
| Race |  |  |  | 0.2390 |
| White or Caucasian | 38 (33%) | 20 (28%) | 18 (25%) |  |
| Hispanic or Latino | 22 (31%) | 16 (23%) | 6 (8%) |  |
| African American | 10 (14%) | 6 (8%) | 4 (6%) |  |
| Asian | 1 (1%) | 0 | 1 (1%) |  |
| BMI |  |  |  | 0.4779 |
| Normal | 16 (23%) | 10 (14%) | 6 (9%) |  |
| Overweight | 21 (30%) | 10 (14%) | 11 (16%) |  |
| Obese | 33 (47%) | 21 (30%) | 12 (17%) |  |
| Dwelling |  |  |  | 0.7017 |
| With family members | 48 (68%) | 27 (38%) | 21 (30%) |  |
| Assisted living | 9 (13%) | 5 (7%) | 4 (6%) |  |
| Alone | 9 (13%) | 6 (9%) | 3 (4%) |  |
| Homeless | 5 (7%) | 4 (6%) | 1 (1%) |  |
| Symptom onset day when blood drawn<br>Mean ± SEM | 11.2 ± 0.92 | 10.2 ± 1.1 | 12.7 ± 1.6 | 0.2010 |
| Diabetes |  |  |  | 0.2258 |
| Yes | 28 (40%) | 19 (27%) | 9 (13%) |  |
| No | 43 (61%) | 23 (32%) | 20 (28%) |  |
| Base line lung disease |  |  |  | 1.0000 |
| Yes<sup>s</sup> | 8 (11%) | 5 (7%) | 4 (6%) |  |
| No | 63 (87%) | 37 (52%) | 25 (35%) |  |
| Respiratory Pathogen Panel |  |  |  | 1.0000 |
| Positive | 6 (9%)<sup>q</sup> | 4 (6%)<sup>y</sup> | 2 (3%) |  |
| Negative | 55 (91%) | 38 (53%) | 27 (38%) |  |
| Absolute lymphocyte count (10<sup>9</sup>/L)<br>Mean ± SEM | 0.71 ± 0.06 | 0.62 ± 0.05 | 0.84 ± 0.14 | 0.1287 |
| D-Dimer (ng/mL) |  |  |  | 0.0289<sup>¥</sup> |
| Median | 850 | 2180 | 431 |  |
| 95% Quantiles | (103, 59640) | (104, 59640) | (103, 2669) |  |
| Mean ± SEM<sup>‡</sup> | 7.0 ± 0.25 | 7.4 ± 0.33 | 6.3 ± 0.27 | 0.0099 |
| Highest level of treatment |  |  |  | 0.1887 |
| Room air | 17 (26%) | 11 (16%) | 7 (10%) |  |
| Nasal cannula | 32 (45%) | 15 (21%) | 17 (24%) |  |
| Intubation | 19 (27%) | 14 (20%) | 5 (7%) |  |
| Extracorporeal membrane oxygenation | 1 (1%) | 1 (1%) | 0 |  |
| Outcome |  |  |  | 0.2277 |
| Deceased | 7 (10%) | 6 (8%) | 1 (1%) |  |
| Survived | 64 (90%) | 36 (51%) | 28 (39%) |  |

§ Asthma and/or chronic obstructive pulmonary disease (COPD) q Positives included: Coronavirus NL63, Coronavirus OC43, Rhinovirus/Enterovirus y Positives included: H1N1, Respiratory syncytial virus A and B ‡ After natural logarithmic transformation ¥ Non-parametric (Wilcoxon) test SEM: Standard error of mean

**Table 2.** Performance comparisons of 4 assays in COVID-19 patients.

| Sample number |  | SQ IgM/IgG<sup>#</sup> |  | Total |
| --- | --- | --- | --- | --- |
|  |  | Pos | Neg |  |
| Wondfo | Pos | 65 (58%) | 6<sup>§</sup> (5.3%) | 71 |
| Total Ab | Neg | 7<sup>q</sup> (6.2%) | 35 (31%) | 42 |
| Total |  | 72 | 41 | 113 |
| % overall agreement (95% CI): 88.5 (81.3, 93.5)<br>% positive percent agreement (95% CI): 90.3 (81.3, 95.2)<br>% negative percent agreement (95% CI): 85.4 (71.6, 93.1)<br>Kappa value (95% CI): 0.75 (0.62-0.88) |  |  |  |  |
| Sample number |  | SQ IgG |  | Total |
|  |  | Pos | Neg |  |
| Abbott IgG | Pos | 58 (52%) | 5<sup>y</sup> (4.5%) | 63 |
|  | Neg | 1<sup>‡</sup> (1%) | 48 (43%) | 49 |
| Total |  | 59 | 53 (42.9) | 112 |
| % overall agreement (95% CI): 94.6 (88.8, 97.5)<br>% positive percent agreement (95% CI): 98.3 (91, 99.7)<br>% negative percent agreement (95% CI): 90.6 (79.7, 95.9)<br>Kappa value (95% CI): 0.89 (0.81-0.98) |  |  |  |  |

#: Either IgM or IgG positive is considered as positive
§: Abbott IgG is positive in 3 of 6; SQ IgG is all negative.

**Table 3.**
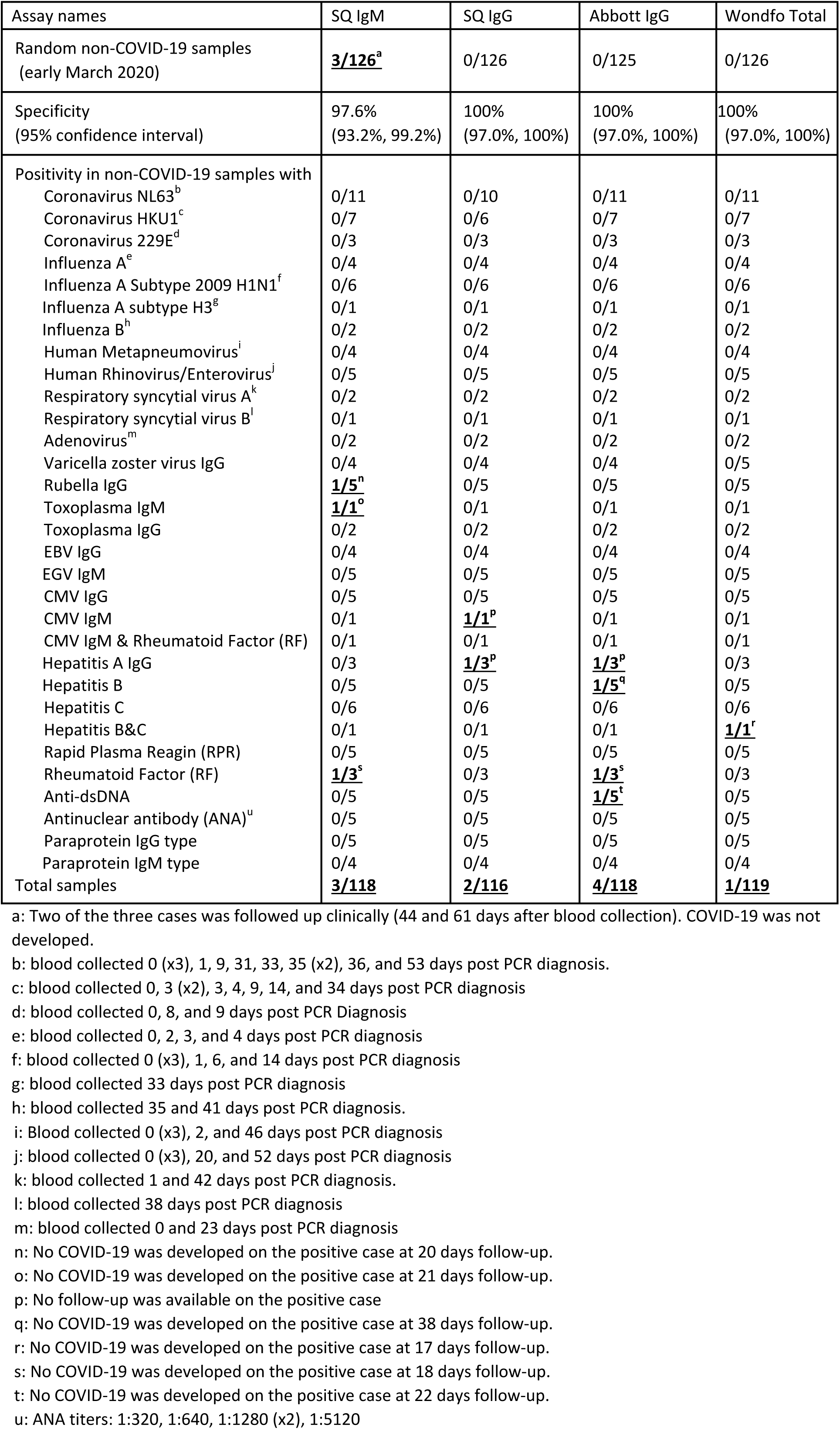
Positive rates of four assays on negative samples and interferences/cross-reactions

| Assay names | SQ IgM | SQ IgG | Abbott IgG | Wondfo Total |
| --- | --- | --- | --- | --- |
| Random non-COVID-19 samples<br>(early March 2020) | **<u>3/126<sup>a</sup></u>** | 0/126 | 0/125 | 0/126 |
| Specificity<br>(95% confidence interval) | 97.6%<br>(93.2%, 99.2%) | 100%<br>(97.0%, 100%) | 100%<br>(97.0%, 100%) | 100%<br>(97.0%, 100%) |
| Positivity in non-COVID-19 samples with |  |  |  |  |
| Coronavirus NL63<sup>b</sup> | 0/11 | 0/10 | 0/11 | 0/11 |
| Coronavirus HKU1<sup>c</sup> | 0/7 | 0/6 | 0/7 | 0/7 |
| Coronavirus 229E<sup>d</sup> | 0/3 | 0/3 | 0/3 | 0/3 |
| Influenza A<sup>e</sup> | 0/4 | 0/4 | 0/4 | 0/4 |
| Influenza A Subtype 2009 H1N1<sup>f</sup> | 0/6 | 0/6 | 0/6 | 0/6 |
| Influenza A subtype H3<sup>g</sup> | 0/1 | 0/1 | 0/1 | 0/1 |
| Influenza B<sup>h</sup> | 0/2 | 0/2 | 0/2 | 0/2 |
| Human Metapneumovirus<sup>i</sup> | 0/4 | 0/4 | 0/4 | 0/4 |
| Human Rhinovirus/Enterovirus<sup>j</sup> | 0/5 | 0/5 | 0/5 | 0/5 |
| Respiratory syncytial virus A<sup>k</sup> | 0/2 | 0/2 | 0/2 | 0/2 |
| Respiratory syncytial virus B<sup>l</sup> | 0/1 | 0/1 | 0/1 | 0/1 |
| Adenovirus<sup>m</sup> | 0/2 | 0/2 | 0/2 | 0/2 |
| Varicella zoster virus IgG | 0/4 | 0/4 | 0/4 | 0/5 |
| Rubella IgG | **<u>1/5<sup>n</sup></u>** | 0/5 | 0/5 | 0/5 |
| Toxoplasma IgM | **<u>1/1<sup>o</sup></u>** | 0/1 | 0/1 | 0/1 |
| Toxoplasma IgG | 0/2 | 0/2 | 0/2 | 0/2 |
| EBV IgG | 0/4 | 0/4 | 0/4 | 0/4 |
| EGV IgM | 0/5 | 0/5 | 0/5 | 0/5 |
| CMV IgG | 0/5 | 0/5 | 0/5 | 0/5 |
| CMV IgM | 0/1 | **<u>1/1<sup>p</sup></u>** | 0/1 | 0/1 |
| CMV IgM & Rheumatoid Factor (RF) | 0/1 | 0/1 | 0/1 | 0/1 |
| Hepatitis A IgG | 0/3 | **<u>1/3<sup>p</sup></u>** | **<u>1/3<sup>p</sup></u>** | 0/3 |
| Hepatitis B | 0/5 | 0/5 | **<u>1/5<sup>q</sup></u>** | 0/5 |
| Hepatitis C | 0/6 | 0/6 | 0/6 | 0/6 |
| Hepatitis B&C | 0/1 | 0/1 | 0/1 | **<u>1/1<sup>r</sup></u>** |
| Rapid Plasma Reagin (RPR) | 0/5 | 0/5 | 0/5 | 0/5 |
| Rheumatoid Factor (RF) | **<u>1/3<sup>s</sup></u>** | 0/3 | **<u>1/3<sup>s</sup></u>** | 0/3 |
| Anti-dsDNA | 0/5 | 0/5 | **<u>1/5<sup>t</sup></u>** | 0/5 |
| Antinuclear antibody (ANA)<sup>u</sup> | 0/5 | 0/5 | 0/5 | 0/5 |
| Paraprotein IgG type | 0/5 | 0/5 | 0/5 | 0/5 |
| Paraprotein IgM type | 0/4 | 0/4 | 0/4 | 0/4 |
| Total samples | **<u>3/118</u>** | **<u>2/116</u>** | **<u>4/118</u>** | **<u>1/119</u>** |

a: Two of the three cases was followed up clinically (44 and 61 days after blood collection). COVID-19 was not developed.
b: blood collected 0 (x3), 1, 9, 31, 33, 35 (x2), 36, and 53 days post PCR diagnosis.
c: blood collected 0, 3 (x2), 3, 4, 9, 14, and 34 days post PCR diagnosis
d: blood collected 0, 8, and 9 days post PCR Diagnosis
e: blood collected 0, 2, 3, and 4 days post PCR diagnosis
f: blood collected 0 (x3), 1, 6, and 14 days post PCR diagnosis
g: blood collected 33 days post PCR diagnosis
h: blood collected 35 and 41 days post PCR diagnosis.
i: Blood collected 0 (x3), 2, and 46 days post PCR diagnosis j: blood collected 0 (x3), 20, and 52 days post PCR diagnosis k: blood collected 1 and 42 days post PCR diagnosis. l: blood collected 38 days post PCR diagnosis m: blood collected 0 and 23 days post PCR diagnosis n: No COVID-19 was developed on the positive case at 20 days follow-up. o: No COVID-19 was developed on the positive case at 21 days follow-up. p: No follow-up was available on the positive case q: No COVID-19 was developed on the positive case at 38 days follow-up. r: No COVID-19 was developed on the positive case at 17 days follow-up. s: No COVID-19 was developed on the positive case at 18 days follow-up. t: No COVID-19 was developed on the positive case at 22 days follow-up. u: ANA titers: 1:320, 1:640, 1:1280 (x2), 1:5120

Of all the 113 samples available from the 71 patients, 105 samples were selected to evaluate antibody positive rates every two days (1-2, 3-4, 5-6, 7-8, 9-10, 11-12, 13-14, and ≥ 15 days post symptom onset). Duplicate samples in the same time frame were not used. Seroconversion date was defined as the middle point between the date of the last negative and the date of the first positive in one patient.

To obtain more precise specificities for SQ IgM, SQ IgG and Abbott IgG, 1063 serum or plasma samples were collected before the pandemic started in the United States (January 2020), including 500 samples originally for reference range determination of a troponin assay, 371 prenatal samples for reference range determination of quadruple tests, 50 pre-pandemic samples from transfusion service and 21 pre-pandemic plasma segments from the Rhode Island Blood Center.

Not all the samples were available for all four tests. Case numbers in the tables and figures may have small difference (up to 4); however, the results and conclusions were not compromised.

### Lateral flow assays

SARS-CoV-2 Total Antibody Test (Wondfo, Guangzhou, China) and STANDARD Q COVID-19 IgM/IgG Duo Test kits (SD Biosensor, Gyeonggi-do, Korea) were purchased from the manufacturers and the assays were performed following the manufacturer’s protocols (8, 9). Briefly, 10 µl of serum or plasma was applied to the designated area of the lateral flow strip following three drops of buffer. Positive result was indicated by a visible band in the designated area accompanied with an appropriate control band. Over 90% of the reading was performed by one investigator (K.J.P.) to ensure consistency.

### Chemiluminescent assay

SARS-CoV-2 IgG test reagents were purchased from the manufacturer (Abbott Diagnostics, Lake Forest, IL). The assays were performed on an Abbott Architect i1000 analyzer following the manufacturer’s protocol. The assay was calibrated initially and with any subsequent reagent lot. A positive and negative Control was run at the start of each batch of antibody testing per manufacturers protocol (10). Serum and plasma samples were both accepted by the assay. Samples with signal-to-cutoff (S/CO) ratio greater than or equal to 1.4 were considered positive.

### Data analysis

The data collected were analyzed on a statistical package, JMP Pro 14.0 (SAS Institute, Cary, NC). Categorical data were analyzed via Chi-square analysis or Fisher’s exact test whenever appropriate. Wilcoxon method was used in parametric test. 95% confidence intervals were calculated for the sensitivity and specificity.

## Results

### Clinicopathologic features of COVID-19 patients in the study

Table 1 summarizes the clinicopathologic features of 71 PCR-confirmed COVID-19 patients in this series, including 42 males and 29 females. Average age of the males was 8.1 years younger than that of the females (P=0.0470). About one third (38) of the patients were White or Caucasian and 31% (22) were Hispanic or Latino. Africa Americans consisted of 14% (10) of the patients. There was one Asian patient. Many of the patients were either overweight (30%) or obese (47%). Most of them 48 (68%) lived with family members, 9 (13%) in assisted facilities, 9 (13%) alone, and 5 (7%) were homeless. Forty percent (40%) of them had diabetes. Only 8 (11%) had baseline respiratory illnesses, mainly asthma and/or chronic obstructive pulmonary disease (COPD). Eight (11%) had positive findings in upper respiratory virus testing (ePlex Respiratory Pathogen Panel), including coronavirus NL63, coronavirus OC43, and rhinovirus/enterovirus, influenza A subtype H1N1, and respiratory syncytial virus A and B. Patients had decreased absolute lymphocyte count at 0.71×10^9^/L on average when the diagnosis of COVID-19 was made. The highest D-Dimer level in the disease course was significantly higher in male patients, with median of 2180 ng/mL, compared to 431 in female patients (P=0.0289; P= 0.0099 after logarithmic transformation). Among the highest level of oxygen requirement in the disease course, 32 (45%) needed oxygen through nasal cannula, 19 (27%) required intubation, 1 needed extracorporeal membrane oxygenation, and the remaining 17 (26%) maintained a satisfactory oxygen saturation on room air. Seven patients died including 6 males and 1 female.

### Antibody detection in early disease stages by four different tests

The patients’ blood samples were collected on average 11.2 days post-symptom onset (Table 1). The samples were grouped every two days within the first 2 weeks starting on the symptom onset day 0, and tested by SQ IgM, SQ IgG, Abbott IgG and Wondfo Total antibody assays. Positive results appeared as early as days 3-4 for SQ IgM, days 5-6 for SQ IgG, days 7-8 for Abbott IgG and Wondfo Total. After 14 days, all the samples were positive by SQ IgG, Abbott IgG and Wondfo Total.

The time points and test results related to seroconversion are listed in Supplemental Table 1. Twenty-three events of seroconversion were recorded and summarized in Figure 2. SQ IgM recorded 7 seroconversions, dated from post-symptom onset days 5.5 to 11, 7.9 days on average. SQ IgG recorded 8 seroconversions, dated from post-symptom onset days 5.5 to 10, 7.6 days on average. Abbott IgG recorded 8 seroconversions, dated from post-symptom onset days 5.5 to 10.5, 7.6 days on average. Wondfo Total Antibodies recorded 10 seroconversions, dating from 5.5 to 11 days, 8.1 days on average. There was no statistical difference among the seroconversion times of all the assays (Figure 2).

**Figure 1.**
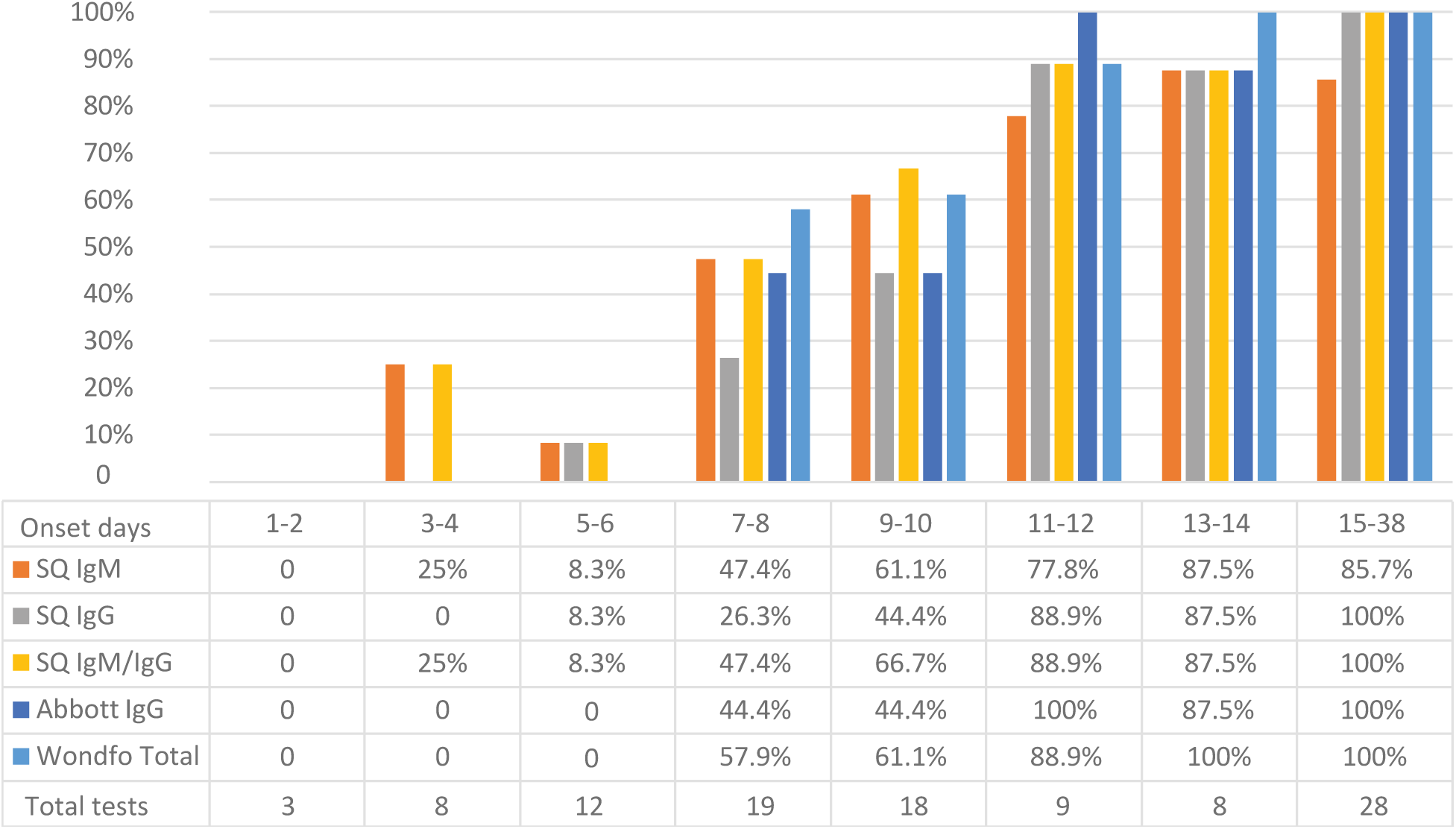
Positive rates of four tests based on symptom onset days. SQ IgM: STANDARD Q COVID-19 IgM (SD BIOSENSOR); SQ IgG: STANDARD Q COVID-19 IgG (SD BIOSENSOR); Wondfo Total: SARS-CoV-2 Total antibody test (Wondfo).

**Figure 2.**
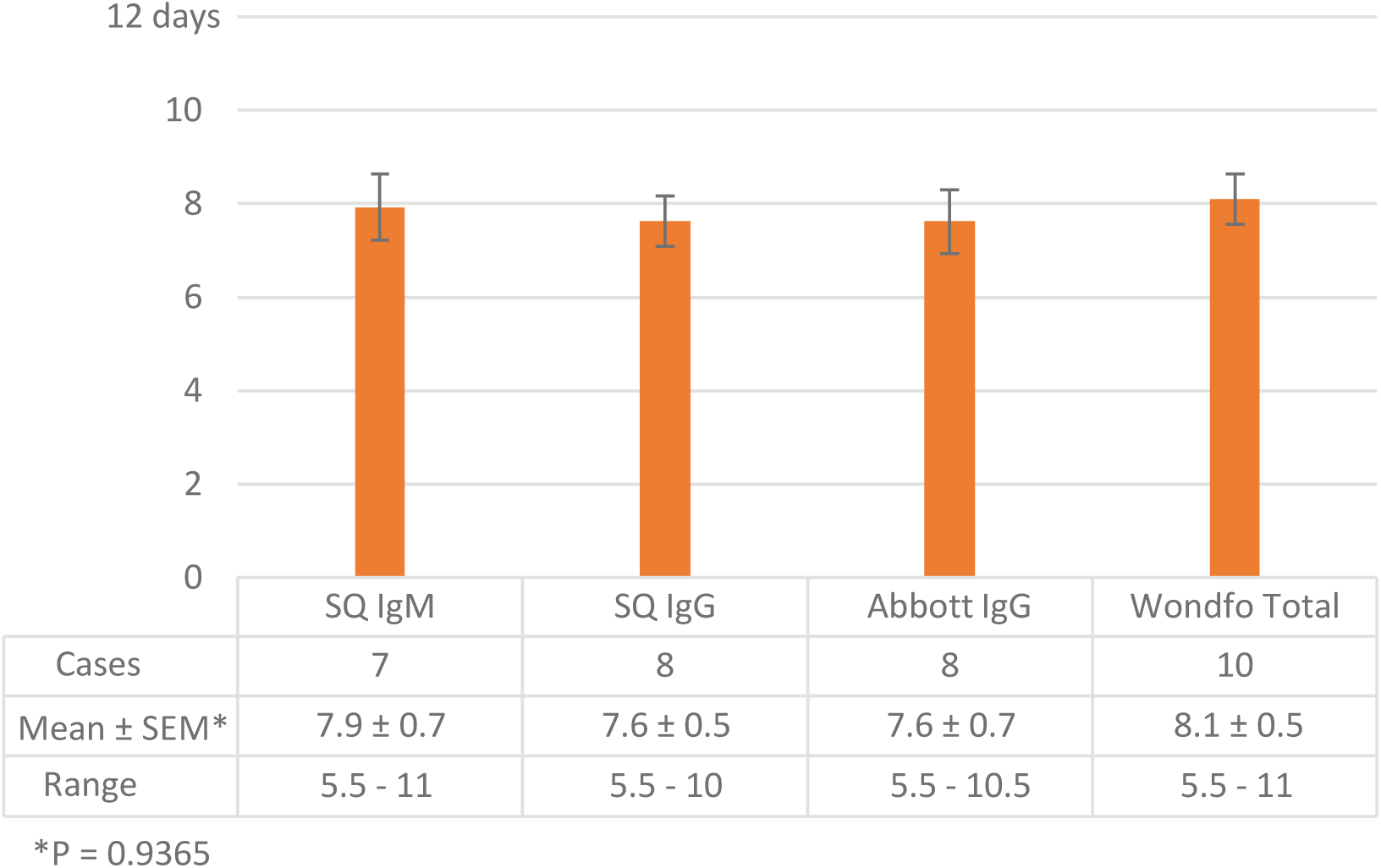
Average seroconversion days detected by four tests. SEM: standard error of mean

### Comparison between IgG assays and IgM/IgG assays

The Wondfo Total Antibody test detects IgG and IgM. Either a positive IgM or a positive IgG will give a positive result. SQ IgM/IgG Duo is packaged as two independent lateral flow devices that are used to assay IgM and IgG in parallel, but their combined interpretation provides a result comparable with that of the Wondfo Total Antibody test. Out of 113 samples from PCR-confirmed patients with antibody results available from all 3 tests, 65 (58%) were positive by Wondfo Total and SQ IgM/IgG and 35 (31%) were negative by all three. Six (5.3%) were positive by Wondfo and negative by SQ IgM and IgG. Seven (6.2%) were negative by Wondfo Total and positive by either of SQ IgM and IgG. The overall agreement was 88.5% and the Kappa value was 0.75. (Table 2)

Between SQ IgG and Abbott IgG, the overall agreement was 94.6% with a Kappa value of 0.89. There were 58 (52%) samples positive by both assays and 48 (43%) negative by both. Five (4.5%) were positive by Abbott IgG and negative by SQ IgG. Of these, 4 were positive by Wondfo Total. One (1%) case was positive by SQ IgG but negative by Abbott IgG and Wondfo Total. (Table 2)

### Cross-reactions, interference and specificities

To obtain the specificities of four tests, 126 assumedly negative samples collected from routine clinical immunology samples were tested. All resulted negative except SQ IgM which had three positives. The specificities and 95% confidence intervals of SQ IgG, Abbott IgG and Wondfo Total were 100% (97.0%, 100%) for all three and 97.6% (93.2%, 99.2%) for SQ IgM based on this series. (Table 3)

To examine the assay’s cross-reactivity to other viruses, the study included 21 samples from patients with seasonal coronavirus NL63 (n=11), HKU1 (n=7) and 229E (n=3). The diagnoses of virus infection were based on nucleic acid testing. No patients with past OC43 infection were evaluated. Blood samples were collected around the diagnoses and after the diagnoses to ensure enough antibody response to be mounted. Eight samples were collected 14 to 53 days after the diagnosis. All 21 samples were negative by the four tests. Similar sample collection scheme was used for other viruses, including influenzas, metapneumovirus, rhinovirus, enterovirus, respiratory syncytial viruses and adenovirus. All these samples were negative by the four tests. (Table 3)

Selected samples with positive IgG and IgM results from other viruses including varicella zoster virus, rubella, Epstein-Barr virus, cytomegalovirus (CMV) and hepatitis viruses were also tested. SQ IgM was positive in one sample with Rubella IgG. SQ IgG was positive in one sample with CMV IgM and one sample with Hepatitis A IgG. Abbott IgG was positive in one sample with Hepatitis A IgG and one sample from an active Hepatitis B patient. Wondfo Total was positive in one sample from a patient with both active Hepatitis B and C. SQ IgM was positive in one sample with Toxoplasma IgM. None of Rapid Plasma Reagin (RPR) samples was positive by any of the tests. (Table 3)

SQ IgM was positive in one sample with Rheumatoid Factor (RF) and Abbott IgG was positive in one sample with RF and one sample with anti-Double Strand DNA (dsDNA). (Table 3) Samples with anti-nuclear antibodies and paraproteins of IgG and IgM types were all negative by four tests. (Table 3)

### False positive rates of SQ IgM, SQ IgG, and Abbott IgG in pre-pandemic samples

Among the 1063 blood samples from frozen pre-pandemic time, SQ IgM had 6 positives from troponin study samples, 2 from plasma segments, and one from prenatal samples. Abbott IgG had 4 positives from troponin study samples. No false positive was seen from SQ IgG results. The specificities and 95% confidence intervals (in parenthesis) for SQ IgM, SQ IgG and Abbott IgG were 98.87% (98.04%, 99.35%), 100% (99.64%, 100%) and 99.62% (99.03%, 99.85%), respectively. (Table 4)

**Table 4.** Positive rate in pre-pandemic and post-pandemic samples for 3 assays

| Assay names | SQ IgM | SQ IgG | Abbott IgG |
| --- | --- | --- | --- |
| Random non-COVID-19 samples<br>(early March 2020) (from Table 3) | **<u>3/126</u>** | 0/126 | 0/125 |
| Pre-pandemic samples from troponin<br>study | **<u>6/500</u>** | 0/500 | **<u>4/498</u>** |
| Pre-pandemic samples from<br>transfusion service | 0/50 | 0/50 | 0/50 |
| Pre-pandemic samples from Rhode<br>Island Blood Center | **<u>2/21</u>** | 0/21 | 0/21 |
| Pre-pandemic samples from prenatal<br>samples | **<u>1/371</u>** | 0/371 | 0/371 |
| Total | **<u>12/1063</u>** | 0/1063 | **<u>4/1059</u>** |
| Specificity<br>(95% confidence interval) | 98.87%<br>(98.04%, 99.35%) | 100%<br>(99.64%, 100%) | 99.62%<br>(99.03%, 99.85%) |

## Discussion

The utility of the COVID-19 serology testing is still subject to debate. As shown in Figure 1, antibodies started to be detected 3-4 days after the symptom onset. Antibody titers continued to increase and after 2 weeks antibodies could be detected by all the tests except SQ IgM whose positive rate peaked at 87.5% around day 13-14. High sensitivity of serology testing after 2 weeks of symptom onset was shown in other studies (11, 12). In one study, 100% sensitivity was seen in Diazyme IgM/IgG assay ≥ 15 days post PCR diagnosis (11) and another study reported 93.8% (95% CI; 82.80-98.69) at ≥14d post symptom onset for Abbott IgG (12).

Serology testing could be used as part of the diagnostic panel after 14 days post symptom onset when the positive rates were the highest and the sensitivity of swab PCR decreased (2, 3). At our institution, it is not uncommon to see patients who were highly suspected of COVID-19 based on signs and symptoms have positive antibody tests while rRT-PCR tests were repeatedly negative. The seroconversion could be detected during the second week post symptom onset (Figure 2). Presence of a seroconversion in a highly suspicious COVID-19 patient should be diagnostic in the right clinical settings. The utility of the testing before 2 weeks post symptom onset should be best decided on a case-by-case basis.

The overall agreement between SQ IgM/IgG and Wondfo Total was as high as 88.5% with Kappa of 0.75. The overall agreement between Abbott IgG and SQ IgG was as high as 94.6% with Kappa of 0.89. The disagreement cases were samples collected during the first 2 weeks of symptoms. Since the tests were generally not recommended before 2 weeks post symptom onset, the difference among the assays would not be clinically significant.

### Specificities and cross-reactions of all four tests

It has been widely considered that the SARS-CoV-2 antibodies may be cross-reactive to seasonal coronaviruses, such as NL63, 229E, HKU1, and OC43. The latter two belong to beta subgroup which also includes SARS-CoV-2. The detection of non-COVID coronaviruses varies from year to year and in some years accounted for as much as 22%-25% of adult respiratory illness (13, 14). Therefore, if cross reactivity did exist, the utility of the SARS-CoV-2 antibody test would be greatly limited. In the current study, among the 21 samples from patients with coronaviruses NL63, HKU1 and 229E, 13 were collected at the time of diagnosis and 8 collected at least 2 weeks after the diagnosis to ensure sufficient development of immune response. All four tests performed well and none of them were reactive to the 21 samples in the study. Consistent with our findings, one study included 5 seasonal coronavirus samples and they were all negative by Abbott IgG (12). Seasonal coronaviruses are known for their short periods of immunity after infection. It is known that antibodies against seasonal coronaviruses reached peak titers in 2 weeks and slowly declined and the protection is largely lost a year later (15); however, due to repeat infection, a report found that adult population had a high seroprevalence of coronaviruses (91.3% for 229E, 59.2% for HKU1, 91.8% for NL63, and 90.8% for OC43) from a United States Metropolitan Population (11). In the current study we included over 1000 samples from pre-pandemic era and overall found very low positive rates in SQ IgM, SQ IgG and Abbott IgG, which echoes the findings from Abbott (10) and an independent study (16). The Wondfo Total was not tested in this evaluation. Given the high prevalence of coronavirus infection in the general population (11), if the cross-reactions were common, the positive rates would be expected to be higher.

The cross-reaction in hepatitis patients was unexpected. Out of 15 samples with hepatitis A, B or C, one sample with Hepatitis A IgG was positive in both the SQ IgG and Abbott tests. One sample with active Hepatitis B was positive by the Abbott IgG test. One sample with active Hepatitis B and C was positive by Wondfo test. It is difficult to determine which hepatitis antibody was indeed cross-reactive because all these samples were expected positive for Hepatitis A IgG and Hepatitis B surface antibodies.

Autoimmune antibodies are known interferences of many antibody tests. SQ IgM and Abbott IgG tests were reactive to a RF positive sample and Abbott IgG test was reactive to an anti-dsDNA positive sample. Special attention should be paid to autoimmune patients when interpreting their positive SARS-COVID-2 antibody results.

### The clinical usefulness of IgM testing

The utility of SARS-COVID-2 IgM testing has not been fully evaluated. Reported specificities of IgM have been suboptimal: only 2 out of 9 tested assays achieved > 95% at the lower end of 95% confidence interval of their specificities (17). However, given the facts that SQ IgM detected IgM only 2 days before SQ IgG detected IgG and that SQ IgM positive rate was only 85.7% in samples over 2 weeks post symptom onset, SQ IgM has marked limit in its clinical utility.

### Use of the antibody testing in community survey

Serologic testing has been used in seroprevalence surveys, including a large-scale geographic survey (18), a community level survey (19), and a special populations survey (20). The key assay characteristic that impacts the accuracy of these surveys is specificity, especially when the disease prevalence is low. It is estimated that for an assay with 99% specificity, the positive predictive value is only ∼50% in a disease with prevalence of 1%. The SQ IgG and Abbott IgG reached over 99% specificities at the lower ends of their 95% confidence intervals. The SQ IgG was negative in all the 1063 negative cases, with 99.64% specificity at the lower end of its 95% confidence interval. Even with 99.64% specificity, the positive predictive value increases to 74% in a disease with prevalence of 1%, and the positive predictive value increases to 93% in a disease with prevalence of 5%.

### Limitation of the study

The main limitation of the study is that the samples from COVID-19 patients were collected from an inpatient population. This group of patients were generally overweight or obese (77%) with high prevalence of diabetes (40%). They tended to have a high D-Dimer levels and marked lymphocytopenia. The clinical symptoms tended to be severe with more being intubated and poor clinical outcomes. The antibody response in this population has been shown to be robust (17). In outpatient population, asymptomatic infected individuals have been reported only with ∼ 10% (28/276) seropositive rates (20). Moreover, asymptomatic and pauci-symptomatic patients could have no detectable antibody response 4 weeks after the diagnosis (21). Another limitation is the limited number of non-COVID positive samples collected at least 2 weeks post symptom onset. More work is needed to assess these tests in this patient population.

Another important question in COVID-19 serology is how long the antibody response will persist. The three samples collected over 30 days post symptom onset had Abbott S/CO reading of 7.58 (31 days), 6.37 (31 days) and 2.43 (35 days). The last one was from a patient with end stage renal disease which is known for its attenuated immune response. The other two were among the most robust immune responses in this cohort (both over 90% quantile of S/CO readings). Obviously, follow-up testing of these patients’ antibody S/CO levels will help to answer the question.

In summary, we validated three SARS-CoV-2 antibody tests, including two lateral flow assays (Wondfo Total Antibody and SQ IgM/IgG combo) and one chemiluminescent assay (Abbott IgG). All tests except SQ IgM performed well with excellent sensitivities two weeks after symptom onset and excellent overall specificities. Hepatitis and autoimmune samples were the main sources of very low interferences/cross-reactions.

## Acknowledgements

The project was funded by Pathology Department of Lifespan Academic Center and Rhode Island Department of Health. The Pathology Department had no role in study design, data collection and interpretation, or the decision to submit the work for publication. Rhode Island Department of Health provided contracted fund to validate Abbott and SD Biosensor assays.

The authors claim no conflict of financial interest related to the project.

The authors want to acknowledge Drs. Angela Caliendo and Jonathon Kurtis to provide valuable feedbacks to the manuscript and the expertise to support the project.

---

**Supplemental Table 1.** Seroconversions detected by four tests.

| Patient ID | Symptom onset | SQ IgM | SQ IgG | Abbott IgG | Wondfo Total |
| --- | --- | --- | --- | --- | --- |
| 17 | Day 4<br>Day 7 | Neg<br>Pos | Neg<br>Pos | Neg<br>Pos | Neg<br>Pos |
| 9 | Day 10<br>Day 12 | Neg<br>Pos | Pos<br>Pos | Pos<br>Pos | Pos<br>Pos |
| 14 | Day 5<br>Day 7 | Pos<br>Pos | Pos<br>Pos | Neg<br>Pos | Neg<br>Neg |
| 19 | Day 10<br>Day 11 | Neg<br>Neg | Neg<br>Neg | Neg<br>Pos | Neg<br>Pos |
| 30 | Day 10<br>Day 12 | Pos<br>Pos | Pos<br>Pos | Pos<br>Pos | Neg<br>Pos |
| 36 | Day 5<br>Day 9 | Neg<br>Pos | Neg<br>Pos | Neg<br>Neg | Neg<br>Pos |
| 42 | Day 9<br>Day 11 | Neg<br>Pos | Neg<br>Pos | Neg<br>Pos | Neg<br>Neg |
| 46 | Day 9<br>Day 13 | Neg<br>Neg | Neg<br>Neg | Neg<br>Neg | Neg<br>Pos |
| 51 | Day 8<br>Day 9 | Pos<br>Pos | Neg<br>Pos | Neg<br>Pos | Pos<br>Pos |
| 69 | Day 5<br>Day 10 | Neg<br>Pos | Neg<br>Pos | Neg<br>Pos | Neg<br>Pos |
| 72 | Day 7<br>Day 8 | Neg<br>Neg | Neg<br>Neg | Neg<br>Neg | Neg<br>Pos |
| 75 | Day 5<br>Day 10 | Neg<br>Pos | Neg<br>Pos | Neg<br>Pos | Neg<br>Pos |
| 78 | Day 4<br>Day 7 | Pos<br>Pos | Neg<br>Pos | Neg<br>Pos | Neg<br>Pos |
| 83 | Day 17<br>Day 25 | Pos<br>Neg | Pos<br>Pos | Pos<br>Pos | Pos<br>Pos |
| 85 | Day 8<br>Day 9 | Neg<br>Neg | Neg<br>Pos | Pos<br>Pos | Pos<br>Pos |
| 86 | Day 12<br>Day 26 | Neg<br>Neg | Pos<br>Pos | Pos<br>Pos | Pos<br>Pos |
| 90 | Day 7<br>Day 9 | Neg<br>Pos | Neg<br>Neg | Neg<br>Neg | Neg<br>Pos |

